# Inferring chromatin looping interactions from microscopy-based measurements: opportunities and pitfalls

**DOI:** 10.64898/2026.11.10.765665

**Authors:** James M. Jusuf, Anders S. Hansen, Christoph Zechner

**Affiliations:** Department of Biological Engineering, Massachusetts Institute of Technology; Cambridge, MA 02139, USA; The Broad Institute of MIT and Harvard; Cambridge, MA 02142, USA; Koch Institute for Integrative Cancer Research; Cambridge, MA 02139, USA; The Novo Nordisk Foundation Center for Genomic Mechanisms of Disease, Broad Institute of MIT and Harvard, Cambridge, MA 02142, USA; Scuola Internazionale Superiore di Studi Avanzati, 34136 Trieste, Italy

**Keywords:** chromatin, chromatin loop, live-cell imaging, super-resolution, microscopy, CTCF, cohesin, enhancer, 3D genome, Bayesian inference, dynamics

## Abstract

Chromatin looping plays essential roles in several nuclear processes and live-cell microscopy has become a key method for studying how chromatin loops behave in time. Extracting dynamical parameters including looping probability, lifetime, and search time from live-imaging data requires inferring looping events from noisy 3D distance trajectories. By analyzing polymer simulation data with known ground truth, we show that with realistic localization precision, the common method of simple thresholding on 3D distance measurements can result in severe misestimation of looping probability, lifetime, and search time. We use simple statistical considerations to show that inference from simple thresholding becomes infeasible when localization error is comparable to the size of the interaction to be studied. Finally, we demonstrate how noise-aware methods can be used to estimate looping parameters more accurately in these regimes.

## Introduction

Chromatin looping regulates key nuclear processes including gene expression, genome integrity, DNA replication, and somatic recombination (*1–7*). Due to the polymeric nature of chromatin, any two loci on the same chromosome encounter each other stochastically at a frequency that decreases with genomic separation (*8*); a loop is characterized by an enrichment of interactions above this expected background (*9*). Loops can be divided into a few major classes with distinct mechanisms and biological roles. First, CTCF– CTCF loops arise when a cohesin complex extruding DNA is stalled by DNA-bound CTCF proteins on both sides (*10–13*). These loops organize the genome into topologically associating domains (TADs) (*14–16*) and play significant roles in gene expression (*17–20*), DNA repair (*21–23*), and recombination (*6, 24*). Second, loops between *cis*-regulatory elements (CREs), including enhancers (E) and promoters (P), are thought to regulate gene expression directly (*1, 2*). Third, loops between genomic regions bound by the Polycomb repressive complex are thought to contribute to the repression of nearby genes (*16, 25–28*). Beyond these, evidence of other types of long-range chromatin interactions has continued to emerge, such as silencer loops (*29–31*), tethering elements (*32*), meta-domains (*33*), range-extenders (*34*), facilitators (*35*), and matchmakers (*36*). Most chromatin loops span genomic separations of tens to hundreds of kilobases (*10, 37–39*), although Polycomb loops can span up to tens of megabases (*25–27*).

Due to their functional importance, chromatin loops have been intensely studied, and numerous methods have been developed to map, study, and quantify loops (*40*). The chromosome conformation capture (3C) family of 3D genomics methods that includes Hi-C (*8, 10, 41*) and Micro-C (*37–39, 42, 43*) has driven major insights into chromatin looping, but carries the major disadvantage of only generating time- and population-averaged snapshots. In comparison, fixed-cell microscopy approaches such as classic DNA-FISH and subsequent FISH-based chromosome tracing methods (*44–47*) allow for single-cell/single-allele resolution, absolute measurements of 3D distances instead of relative sequencing counts, and the ability to multiplex DNA readouts with e.g., nascent transcription or protein localization measurements (*44*). However, fixed-cell microscopy is still limited to single snapshots in time.

In contrast, live-cell microscopy can obtain single-allele measurements at multiple time points while avoiding experimental artifacts caused by cell fixation. Therefore, live-cell methods have emerged as key to studying chromatin looping (*48*), especially in light of increasing evidence that chromatin loops are highly dynamic over seconds to minutes timescales (*49–54*). This is consistent with the current understanding of the dynamic nature of the chromatin polymer (*55–57*) and the kinetics of related processes such as transcription bursting (*53, 58–60*), transcription factor binding (*61*), and DNA doublestrand break repair (*21, 62*). Therefore, precisely measuring the dynamical parameters of chromatin looping is crucial towards understanding chromatin looping mechanisms and how loops interface with other biological processes. We identify three key parameters of interest: the probability of loop formation, the lifetime of looping events, and the search time required for two loci to encounter each other and form a loop (*9, 49–52, 56, 63–65*).

Here, we focus on the analytical challenges of quantifying these dynamical parameters from live-imaging data. We begin by discussing 3D distance as a commonly used indicator of looping events. Using simulated data, we demonstrate that thresholding on 3D distance alone, despite being a simple and intuitive approach, cannot be used to accurately quantify chromatin looping parameters in the presence of realistic localization noise. We then show that the failure of thresholding can be explained using simple statistical considerations. Finally, we discuss how noise-aware Bayesian approaches can be used to overcome these challenges.

### 3D distance as an indicator of looping

Since chromatin constantly undergoes stochastic motion, the two DNA loci that participate in a chromatin loop may exist in a “looped” or “unlooped” configuration at any given time (Fig. **1A**). In fixed- or live-cell microscopy studies, the 3D distance between the two loci typically serves as a readout of the loop’s status: a small 3D distance indicates a looped configuration whereas larger 3D distances indicate an unlooped configuration (Fig. **1A**). The distinction between these two configurations can also be seen in the bimodality of the 3D distance histogram (Fig. **1A**) without measurement noise. Historically, chromosome tracing of *Drosophila* polytene chromosomes was reported as early as 1984 (*66, 67*), and pairwise fluorescence 3D distance measurements were established in 1989 (*68*). These techniques have seen rapid improvements in recent years enabling both high-resolution fixed- and live-cell imaging of pairwise distances. Excellent discussions of these technologies can be found in recent reviews (*44, 48, 57, 69*). However, there remain numerous sources of uncertainty which contribute to unavoidable errors in 3D distance measurements.

**Figure 1:**
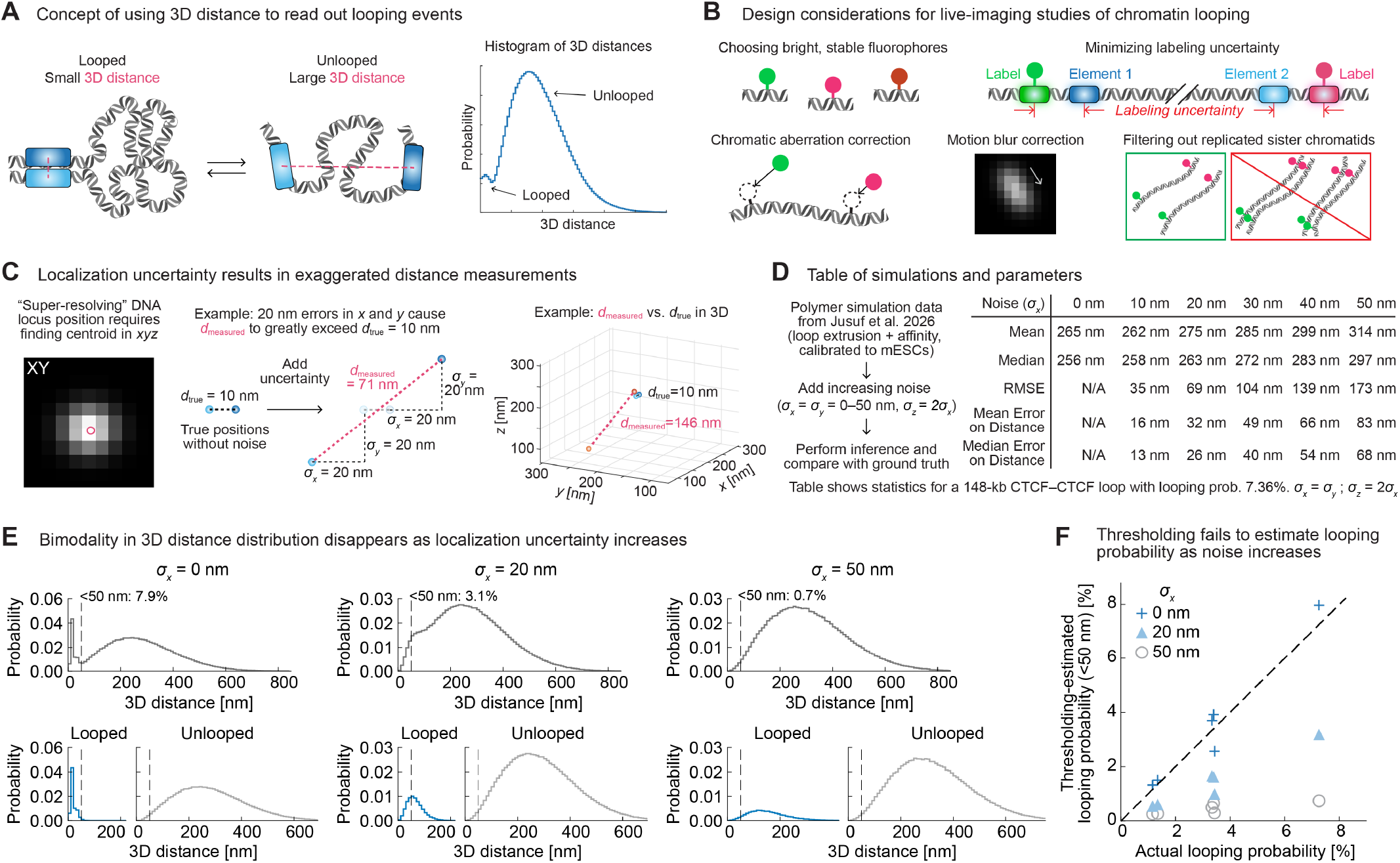
Illustrations of how localization uncertainty will cause measured 3D distances to be substantially larger than true 3D distances, which complicates looping analyses. (A) Illustration of the concept of using 3D distance as a readout of looping. Since when a loop is present between two distal sites the 3D distance ought to be small, we expect a bimodal histogram as shown on the right. (B) Practical considerations for fluorescent labeling of genomic loci besides optical limitations. (C) Illustrations of how optical localization uncertainty corrupts 3D distance measurements. Left: Simulation of “super-resolving” localization. A point source was convolved with a Gaussian point spread function (PSF) and pixelated to simulate an image of a diffraction limited fluorescent DNA locus. The coordinates of the locus are then obtained either using PSF-fitting (*70*) or FracShift (*71*). Because the localization error/uncertainty scales with the inverse square root of the number of detected photons (*70*), reducing *σ*_*x*_ below 20 nm is often quite challenging, especially in live-cell imaging applications where photobleaching must also be minimized. Middle: 2D example where the true distance between the blue dots is 10 nm, but after adding 20 nm error in both *x* and *y*, the measured 2D distance increases to 71 nm. Right: In 3D, this gets even worse since typically *σ*_*z*_ ∼2*σ*_*x*_ and in this instance a true distance of 10 nm (blue dots) can be erroneously measured as 146 nm. (D) In this study, we used realistic 3D polymer simulations from Jusuf et al. 2026 (*9*), re-run with higher time sampling. These simulations include both loop extrusion and CTCF barriers (to make CTCF loops/dots) and affinity-mediated interactions (to make CRE loops/dots). The simulations were calibrated to mESC dynamics (*49*) corresponding to ∼20 ms and 1 kb/21 nm coarse-graining in time and space. See also Fig. 1d in Jusuf et al. 2026 (*9*). The table shows for one CTCF–CTCF loop from Jusuf et al. 2026 (*9*) (148-kb loop (1604–1752 kb) with looping probability 7.36%), the mean and median distance, the Root-Mean Squared Error 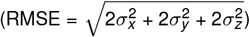, and the error on the 3D distance (⟨|*d*_true_ − *d*_measured_|⟩), for simulations with *σ*_*x*_ = [0, 10, 20, 30, 40, 50] nm noise. (E) Example of 3D distance distribution, introduced in (A), changing from bimodal to unimodal as localization uncertainty increases. The distribution shown is for the 148-kb CTCF–CTCF loop from (D). The “looped” histogram contains all time points where cohesin(s) were observed to connect the loop anchors in the simulation; the “unlooped” histogram contains all other time points. Dashed lines indicate 50-nm threshold, and percentages of distributions below 50 nm are displayed; compare with actual looping probability of 7.36%. (F) Scatterplot of fraction of time below 50-nm threshold versus actual looping probability at three different noise levels (*σ*_*x*_ = 0, 20, and 50 nm). Data points correspond to the 6 different loops analyzed (3 CTCF–CTCF and 3 CRE–CRE). Dashed line indicates a perfect correspondence between fraction of time spent below threshold and actual looping probability.

### Localization uncertainty is unavoidable and causes over-estimation of 3D distances

When designing a chromatin locus-tracking experiment, various known issues should be accounted for in order to minimize localization uncertainty (Fig. **1B**). Fluorescent labels should be as bright and as photostable as possible, as small as possible, and as close to the genomic element of interest (e.g., an enhancer or promoter) as possible without perturbing the function; the elements themselves should also be as small as possible (Fig. **1B**). Even so, labeling uncertainty can easily reach ∼5–25 kb, which corresponds to ∼30–150 nm of localization uncertainty. These labeling considerations have been discussed in previous reviews, along with other important issues including chromatic aberration corrections, filtering of replicated sister chromatids, motion blur, etc. (*1, 48*) (Fig. **1B**). Therefore, we ignore them and model only optical localization uncertainty in our analyses, noting that our results represent an optimistic assessment of true noise conditions.

To illustrate the effect of optical localization uncertainty on 3D distance measurements, we begin by discussing how the position of a single DNA locus is determined in superresolution microscopy. The locus appears as a pixelated, blurred dot in the resulting fluorescence image (Fig. **1C**; the point spread function (PSF) of the microscope describes how a point source will be blurred). The precise *XYZ* coordinates of the locus can then be obtained using methods such as PSF fitting (*70*), FracShift (*71*), or Spotiflow (*72*). In this process, the localization uncertainty *σ* in each coordinate can be modeled as Gaussian and scales with the inverse square root of the number of detected photons (*70*). Live imaging typically sees *σ* ≥20 nm in the *x* - and *y* -dimensions each, whereas fixed-imaging has achieved sub-nanometer precision (*73*). These errors cause a systematic bias towards overestimating the pairwise distance between two loci. For example, 20 nm of localization error in *x* and *y* can cause two loci separated by 10 nm to appear 71 nm apart, a 7-fold overestimation (Fig. **1C**). In 3D, this problem is exacerbated because the axial localization uncertainty is typically ∼2-fold worse (*σ*_*z*_∼ 2*σ*_*x*_ ) and because a 3D distance measurement requires estimating six coordinates: (*x*_1_, *y*_1_, *z*_1_, *x*_2_, *y*_2_, *z*_2_). For example, 30-nm errors in *x* and *y* and 60-nm error in *z* can cause two loci separated by 10 nm to appear 146 nm apart, a 15-fold overestimation (Fig. **1C**). As discussed previously (*1,48,51,65*), measured 3D distances of 100–300 nm therefore should not be interpreted literally and in fact remain consistent with molecular contact at smaller scales.

To quantitatively analyze the effect of localization uncertainty on the inference of looping parameters, we turned to existing simulations of a realistic chromatin polymer model containing cohesin-mediated loop extrusion, CTCF sites, and high-affinity sites (modeling enhancers and promoters) from our previous work (*9*). The usage of simulated data allows for localization error to be added artificially and subsequent inference results compared to their ground-truth values. In the discussions that follow, we analyze 3 CTCF–CTCF loops and 3 affinity-mediated CRE–CRE loops with localization errors ranging from 10 to 50 nm (Fig. **1D**). Considering basic statistics on a 148-kb CTCF–CTCF loop, the mean and median apparent 3D distances increase with increasing noise, as do the RMSE and the errors on those distances (Fig. **1D**). This table also illustrates a challenge in comparing experimental studies that report localization uncertainty using e.g., RMSE (*49*) vs. median error on the distance (*74*) (Fig. **1D**).

### Localization uncertainty obscures the distinction between looped and unlooped configurations

As localization uncertainty increases, the 3D distance distribution changes from bimodal to unimodal; this effect is shown for the aforementioned 148-kb loop in Fig. **1E**. In the absence of noise, nearly all looping events fall within a clear peak below a 50-nm threshold, whereas the 3D distance mostly exceeds this threshold for unlooped configurations. However, even with a modest 20-nm localization uncertainty, the apparent 3D distance in the looped state increases enough that the majority of the looped state appears above the 50-nm threshold. With 50 nm of localization uncertainty, almost all of the looped state appears above the 50-nm threshold (Fig. **1E**). The presence of noise therefore causes an overlap in the 3D distance distributions of the looped and unlooped states, such that a threshold can no longer distinguish between them.

### Localization uncertainty generally causes thresholding to underestimate looping probabilities

Because localization uncertainty causes overestimation of 3D distances (*48*), it generally causes underestimation of looping probabilities when inferred using thresholding. For CTCF– CTCF loops, we define looping probability as the fraction of time that the loop exists in the looped state, characterized by stalled cohesin(s) connecting the anchors of the loop. For affinity-mediated loops, we define looping probability as the fraction of time that the 3D distance is below 50 nm, minus the background probability (calculated from another simulation where the loop anchors have zero affinity). As localization uncertainty increases, the looping probability inferred from thresholding decreases, deviating further from the true value (Fig. **1F**). For example, the aforementioned 148-kb CTCF– CTCF loop has a true looping probability of 7.4%, which is estimated to be 7.9% when thresholding with 0 nm noise. However, the apparent looping probability is only 3.1% with 20 nm noise and 0.7% with 50 nm noise (Figs. **1E** and **1F**). Note that if the looped state is very rare to begin with, thresholding could in principle cause overestimation of looping probability due to false positives from noise, but we did not observe this for any of our loops, whose probabilities range from 1.2% to 7.4%.

### Inferring dynamical parameters of looping in the face of localization uncertainty

Another question of great biological significance is how long chromatin looping interactions last and how often they occur. In this section, we use simulation data to estimate loop lifetimes and search times via simple thresholding: the loci are inferred to be looped at all time points where the 3D distance is ≤ 50 nm and unlooped otherwise. We demonstrate that this approach fails to accurately recover these dynamical parameters when even modest localization uncertainty is introduced.

### Estimating the lifetime of looping events

Due to the different mechanisms underlying CTCF–CTCF and CRE–CRE loops, we consider the estimation of loop lifetimes for each type of loop separately. For CTCF–CTCF loops, the lifetime of a looping event is simply the amount of time that the loop anchors are bridged by one or more stalled cohesin(s) (Fig. **2A**); the mean lifetimes of the three simulated CTCF– CTCF loops in our simulations range from 8–19 minutes in line with recent estimates from live-cell imaging (*49–52*). In live-imaging data, one can define looping events as the periods when the 3D distance is below 50 nm (*74*). These intervals roughly correspond to true looping events in the absence of noise, but the correspondence largely vanishes once noise is introduced (Fig. **2B**; this visualization was inspired by Fig. 2A in Ubertini et al. (*65*), which similarly illustrates the issues with simple thresholding). This is because localization uncertainty introduces single-frame false positive events and also breaks up true sustained contacts into shorter fragments (Fig. **2B**). For example, localization uncertainty of 20 nm can cause the measured 3D distance to exceed 50 nm several times during an 18-minute looping event, making it appear as several 1– 2-minute looping events in close succession. Note that even thresholding with 0 nm noise does not perfectly reflect true looping events because thresholding cannot differentiate between cohesin-mediated loops and proximity due to random polymer motion. Another key observation is that mean estimated loop lifetimes are largely a function of the time between frames, with the mean lifetime approaching the time between frames exactly as localization noise increases (Fig. **2C**). This can cause severe errors in lifetime estimates: for example, with acquisition at Δ*t* = 0.02 s and *σ*_*x*_ = 10 nm noise (representing a spatiotemporal resolution that is vastly better than what is currently experimentally possible), the CTCF loop lifetime is underestimated by approximately 10,000-fold.

**Figure 2:**
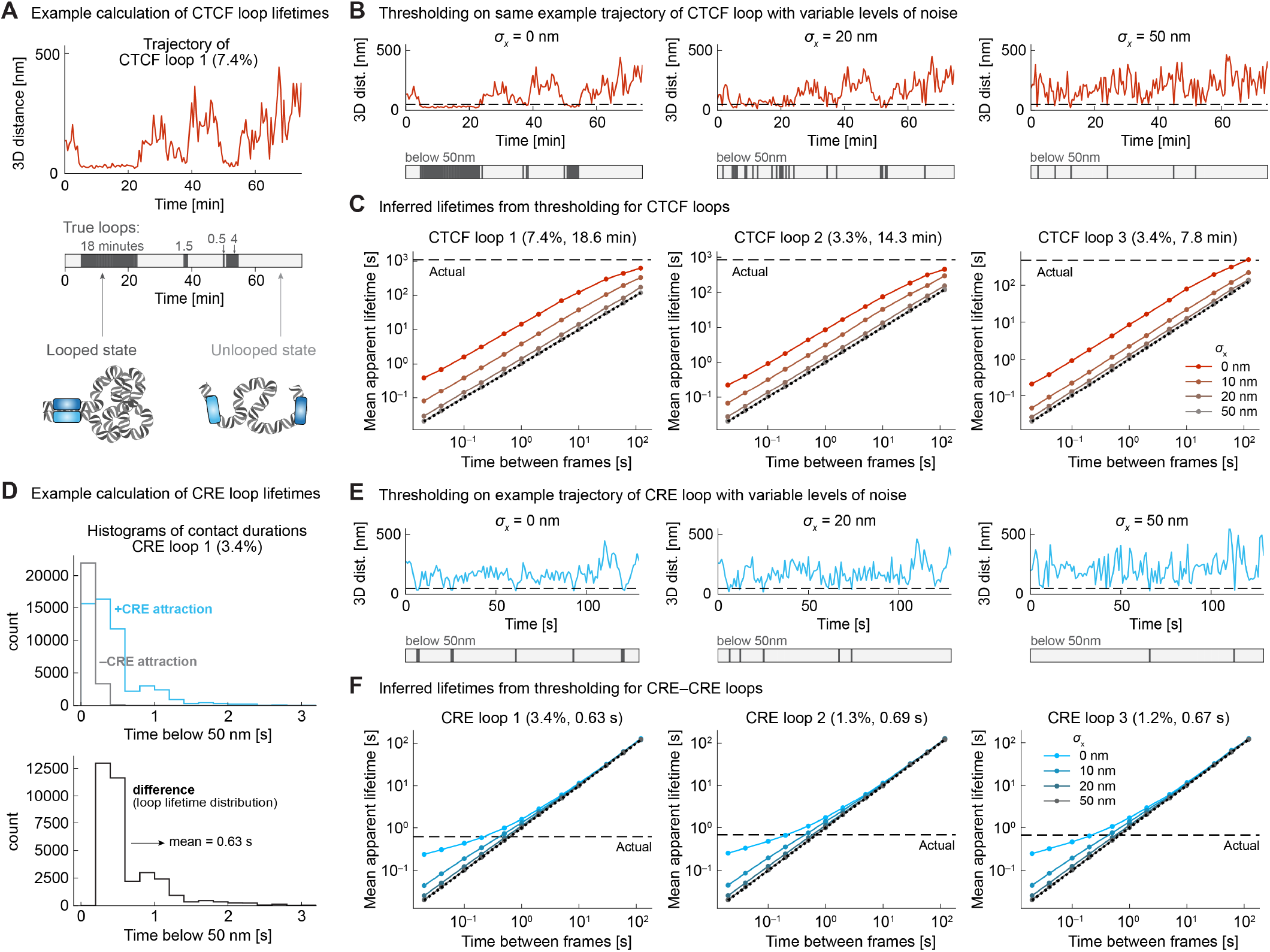
Loop lifetimes cannot be accurately estimated with a simple thresholding approach. (A) Example segment of a 3D distance trajectory of a simulated CTCF–CTCF loop, with true looping events shown below in dark gray, labeled with their lifetimes in minutes. True looping events are defined by presence of cohesin(s) connecting the loop anchors. Time between frames (Δ*t* ) = 30 seconds. (B) The same segment of the 3D distance trajectory shown in (A) with three different noise levels (0, 20, and 50 nm). Dashed line indicates 50-nm threshold, and time points with 3D distance below this threshold are indicated below in dark gray. Δ*t* = 30 seconds. The illustrations used here were inspired by Figure 2A in Ubertini et al. (*65*). (C) For all three simulated CTCF–CTCF loops, mean apparent lifetimes (defined as durations of continuous intervals with apparent 3D distance ≤ 50 nm) are shown as a function of the time between frames Δ*t*, for four different noise levels. Dashed lines indicate true loop lifetimes, and dotted lines indicate mean apparent lifetime = time between frames. (D) Histogram of contact lifetimes for a CRE–CRE loop simulated with attraction (blue) and without attraction (gray). The simulation without attraction is used to establish the baseline of contacts due to random polymer motion. Below is the difference between the distributions, which we define as the loop lifetime distribution. (E) A segment of the 3D distance trajectory of the loop analyzed in (D) with three different noise levels (0, 20, and 50 nm). Dashed line indicates 50-nm threshold, and time points with 3D distance below this threshold are indicated below in dark gray. Δ*t* = 1 second. (F) For all three simulated CRE–CRE loops, mean apparent lifetimes are shown as a function of the time between frames Δ*t*, for four different noise levels. Dashed lines indicate true loop lifetimes, and dotted lines indicate mean apparent lifetime = time between frames.

For CRE–CRE loops, which are modeled as affinity-mediated interactions in our simulation, true looping events are more difficult to define because functional details such as TF-mediated interactions and transcription activation are not explicitly modeled in our simulation. Here, we define the ground-truth CRE–CRE loop lifetime distribution by taking the distribution of intervals with true 3D distance below 50 nm and then subtracting the corresponding distribution in a zero-affinity control condition at Δ*t* = 0.1 s (Fig. **2D**). When inferring lifetimes from thresholding on simulated live-imaging tracks, localization uncertainty once again causes the inferred events to be dominated by single-frame false positives or fragmented true positives (Fig. **2E**). Plotting mean estimated loop lifetimes as a function of localization uncertainty and the time between frames, we observe a similar trend as with the CTCF– CTCF loops, where the mean apparent loop lifetimes approach the time between frames as localization uncertainty increases (Fig. **2F**).

Increasing the threshold or enforcing a minimum duration for inferred contacts may appear to compensate for thresholding artifacts in the presence of localization error; however, these strategies do not circumvent the fundamental issues of thresholding and generate results that are highly dependent on the chosen parameters. To illustrate this, we performed simple thresholding on our simulated chromatin loops with a 200-nm threshold (*52, 75*) and found that the inferred lifetimes once again did not reflect their true values, and were largely a function of the time between frames (Fig. **S1**). We then analyzed the effect of setting a temporal threshold in addition to the 200-nm spatial threshold. Following a previously described strategy (*52,76*), we set a minimum contact time of *T* frames, identified intervals for which the 3D distance was below 200 nm for at least *T* frames, and calculated a moving average with a window size of *T* frames. Consistent with previous results (*52*), we found that this thresholding scheme could reproduce realistic values for loop lifetimes for a specific parameter set (e.g., Δ*t* = 30 s, *T* = 6 frames, and *σ*_*x*_ = 20 nm result in mean inferred lifetimes of 9–12 min, similar to the true values of 8–19 min), but the exact result varied greatly with Δ*t* and the chosen value of *T* (Fig. **S2A**). This trend existed regardless of the amount of localization error present, but higher localization error generally resulted in lower apparent lifetimes (Fig. **S2A**). We observed a similar result for our simulated CRE–CRE loops (Fig. **S2B**); since these events were so short-lived, low sampling frequencies (e.g., Δ*t* = 30 s) resulted in major overestimation of the lifetime.

In summary, apparent loop lifetimes from thresholding are primarily a reflection of the acquisition parameters of the live-imaging experiment (frame rate set on the microscope) rather than the underlying biology (chromatin looping event lifetime). Indeed, in one study where a simple thresholding approach was applied to live-cell trajectories, the thresholding-determined contact lifetime of 0.6 seconds almost exactly matches the acquisition timestep of 0.5 seconds (*74*) as expected from Figs. **2C** and **2F**. Another important consequence is that the effect of perturbation experiments on loop lifetimes may be obscured, since the apparent lifetime estimate is dominated by false positives from noise, regardless of the frequency and lifetime of true looping events.

### Estimating the search time for loop formation

Complementary to loop lifetime measurements, live-imaging experiments can also be used to measure the search time associated with loop formation. Here, we define the search time as the amount of time for a chromatin loop to transition from an unlooped state to a looped state. In the following section, we describe three methods that have been used to estimate search times: direct observation in inducible loops, simulation-based prediction constrained by locus tracking data, and thresholding-based estimation on live-imaging trajectories.

The Chemically Induced Chromosomal Interaction (CICI) experimental platform (*63*) in budding yeast has enabled chromatin search events to be observed directly with imaging (*64*). In this system, a LacO array and a TetO array are inserted into two genomic positions, and fusion fluorophores LacI-GFP and TetR-mCherry are expressed, allowing the two arrays to be localized in 3D using super-resolution live-cell fluorescence microscopy. In addition, LacI-FKBP12 and TetR-FRB fusion proteins are expressed and bind to the arrays, such that the addition of rapamycin causes FKBP12 and FRB to bind strongly and effectively irreversibly, inducing a highly stable chromatin loop (Fig. **3A**). Here, the search time is the amount of time from the addition of rapamycin until the distance between the arrays reaches a certain threshold, also known as the CICI formation time (CFT) (Fig. **3A**). Measuring the search times for a variety of array positions with different separations can be used to empirically determine a general relationship between search time and the distance between the loop anchors (Fig. **3A**). A major advantage of CICI is that it allows direct experimental observation of first passage times and is compatible with perturbation experiments. A disadvantage is the need to insert LacO and TetO arrays and carefully optimize the expression of the LacI-FKBP12 and TetR-FRB fusion proteins (*63, 64*).

**Figure 3:**
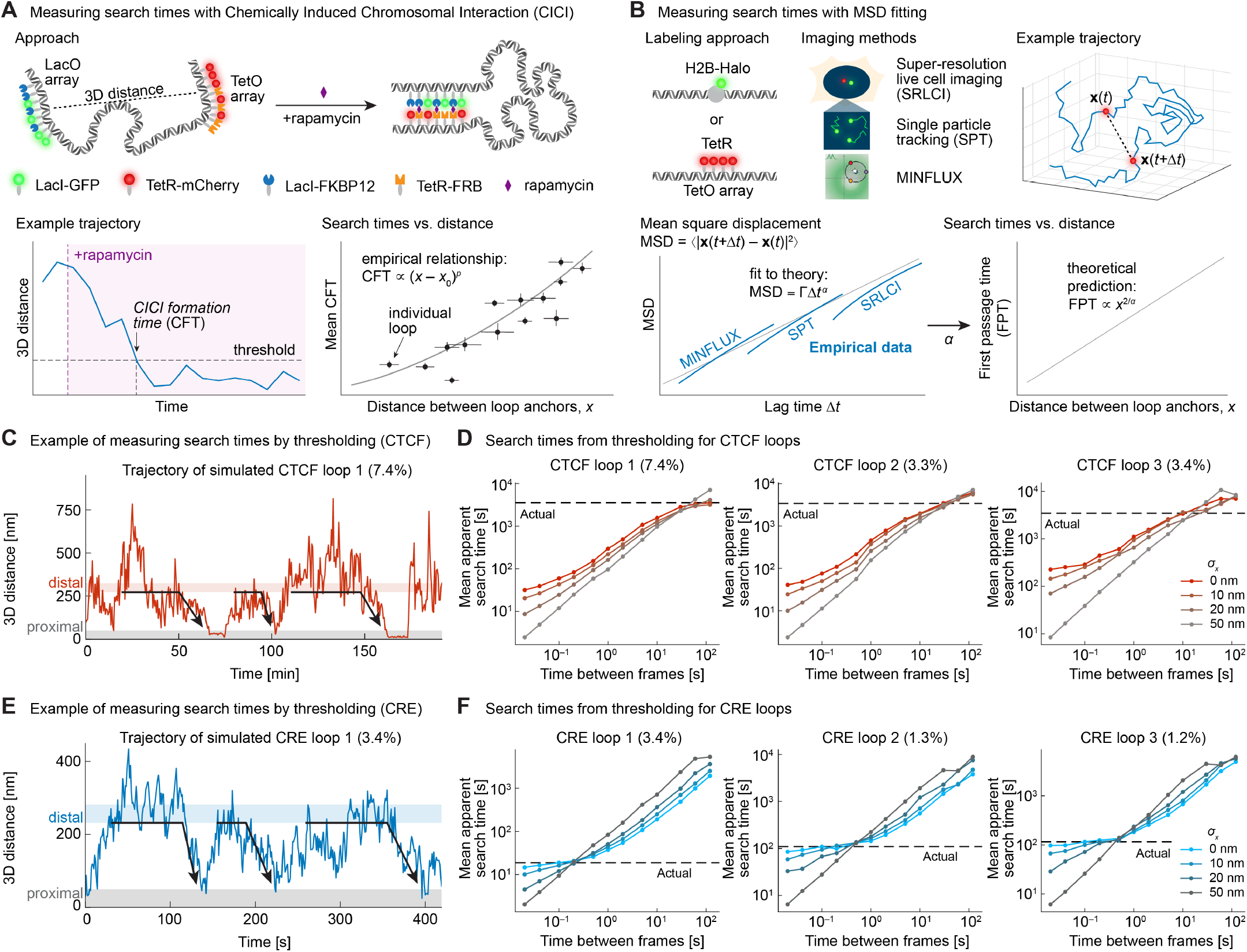
Using thresholding to estimate search times can yield highly variable results depending on localization uncertainty and acquisition parameters. (A) Overview of how Chemically Induced Chromosomal Interaction (CICI) can be used to measure search times of chromatin loops. In this system, a chromatin loop can be induced with the addition of rapamycin and observed using fluorescent probes. An example trajectory of 3D distance vs. time is shown, and the search time (called the CICI formation time (CFT)) is indicated. A plot with hypothetical observations for mean CFT vs. distance between loop anchors is shown, illustrating the relationship between the two quantities that can be fitted to a power law. (B) Overview of how locus tracking data can be used to estimate search times of chromatin loops. First, chromatin is labeled with sparse H2B labeling or at a specific TetO array. Then, super-resolution live-cell imaging (SRLCI), single particle tracking (SPT), or MINFLUX imaging is used to obtain 3D trajectories of the tracked loci. The empirical MSDs from these trajectories can be fitted to a theoretical model, which then predicts a power-law relationship between the first passage time (search time) and the distance between the loop anchors. (C) Example segment from a 3D distance trajectory of a simulated CTCF–CTCF loop. The proximal state (0–50 nm) is highlighted in gray, and the distal state (median to median+50 nm) is highlighted in red. For each entry into the proximal state, the search time is defined as the average time from the preceding distal states to this entry time. Time between frames Δ*t* = 30 seconds. (D) For all three simulated CTCF–CTCF loops, mean apparent search times are shown as a function of the time between frames Δ*t*, for four different noise levels. Dashed lines indicate true mean search times; true search times are defined as in (C) but taking true CTCF–CTCF looping events to define the proximal state instead of 3D distance. (E) Example segment from a 3D distance trajectory of a simulated CRE–CRE loop. The proximal state (0–50 nm) is highlighted in gray, and the distal state (median to median+50 nm) is highlighted in blue. For each entry into the proximal state, the search time is defined as the average time between the preceding distal states to this entry time. Time between frames Δ*t* = 1 s. (F) For all three simulated CRE–CRE loops, mean apparent search times are shown as a function of the time between frames Δ*t*, for four different noise levels. Dashed lines indicate true mean search times; true search times are defined as in (E), specifically with 0 nm noise when sampling at Δ*t* = 0.1 s.

A second approach that can be used to estimate the search times associated with chromatin interactions is to fit histone or locus tracking data, estimate the subdiffusive exponent and prefactor, and then use these experimental constraints to simulate chromatin dynamics in the absence of noise and estimate search times from simulated trajectories (*56, 65, 77–82*). Typically, either histone proteins or specific loci are labeled and tracked over time in such studies (Fig. **3B**). Given a particular MSD exponent of *α*, theory predicts that search time (defined as first passage time) scales as *x* ^2*/α*^, where *x* is the distance between the loop anchors (*56*). An advantage of this approach is that it is relatively straightforward and does not require genome-editing. However, there are also several disadvantages. First, the search time is not directly measured but indirectly estimated. Second, one must assume an underlying model (e.g., fractional Brownian motion) and if the modeling assumptions are not correct, the estimated search times also may not be correct. Third, it is challenging to obtain data over a sufficient range of timescales to rigorously fit the subdiffusive exponent (*83*), though this was recently achieved by integration of MINFLUX, single-particle tracking (SPT), and super-resolution live-cell imaging (SRLCI) (Fig. **3B**) which enabled tracking across seven orders of magnitude in time (*56*).

Finally, we revisit our polymer simulations to investigate whether search times can be inferred by thresholding on 3D trajectories from two-locus live imaging. We apply the definition of search times used in one recent live-imaging study (*74*): the average amount of time to transition from a distal state to a proximal state. The proximal state is defined as the intervals for which the distance between the loop anchors is below a threshold of 50 nm, and the distal state is defined as a 3D distance between *m* and *m* + 50 nm, where *m* is the median 3D distance. An example of search times overlaid with the trajectory of the 148-kb simulated CTCF–CTCF loop is shown in Fig. **3C**. However, when investigating the general trend across three simulated CTCF–CTCF loops, for different values of the time between frames and localization error, we find that the apparent search time strongly depends on both quantities (Fig. **3D**). Similarly, when using thresholding to estimate search times for three simulated CRE–CRE loops, the apparent search time once again depends strongly on the time between frames and localization error (Figs. **3E** and **3F**). Interestingly, in the absence of localization error, thresholding can accurately estimate search times if sampling is sufficiently frequent; this is indicated by a plateau of the *σ*_*x*_ = 0 nm curve below Δ*t* = 0.2 s (Fig. **3F**). However, in the regime of realistic noise for super-resolution imaging (*σ*_*x*_≥ 20 nm), search time estimates from simple thresholding are highly affected by noise-induced artifacts in a frame rate-dependent manner, and thus reflect the microscope acquisition settings instead of the biological system.

### Statistical considerations for inferring proximity from 3D distance trajectories

Thus far, we have investigated the use of simple thresholding to quantify looping probability (Fig. **1**), lifetime (Fig. **2**), or search times (Fig. **3**) by comparing inference results under realistic localization error against known ground-truth. In all cases, the estimates from simple thresholding deviate dramatically from the true values once localization error is introduced. Next, we show that these results can be explained merely from statistical considerations, and we derive theoretical limits that quantify how well simple thresholding can possibly perform under different noise levels. Then, we demonstrate a noiseaware Bayesian thresholding method as an alternative to simple thresholding that infers looping events more accurately in the face of localization uncertainty.

### Mathematical formalization of inference problem

We consider a distance trajectory sampled at discrete time points *t*_1_, …, *t*_*N*_ which we denote by *V* = (*V*_1_, …, *V*_*N*_ ), where 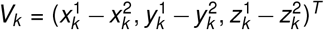 is a 3D distance vector at time point *t*_*k*_ . For simplicity, we consider equidistant sampling times with measurement intervals *t*_*k*+1_− *t*_*k*_ = Δ*t* for all *k* . Moreover, the individual dimensions of *V*_*k*_ are considered to be statistically independent. To define proximity, we segment the trajectory *V* into regions where *d* (*V*_*k*_ ) ≤*α*, where *d* is a metric on ℝ^3^ such as the Euclidean distance and *α* is the proximity threshold. We denote the proximity trajectory as *U* = (*u*_1_, …, *u*_*N*_ ) where *u*_*k*_ = 1 if *d* (*V*_*k*_ ) ≤*α* and *u*_*k*_ = 0 otherwise. The proximity trajectory *U* contains information about the frequency and lifetime of proximity events. We refer to the experimentally measured distance trajectory as *W* = (*W*_1_, …, *W*_*N*_ ), where *W*_*k*_ = *V*_*k*_ + *ϵ*_*k*_ with *ϵ*_*k*_ ∼*N*(0, Σ_*ϵ*_) as Gaussian noise vectors that are independent across time points with covariance Σ_*ϵ*_ = 2 · diag 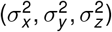. The factor of 2 arises because each distance is computed as a difference between two positions. For simplicity, we consider isotropic distance error in the following such that 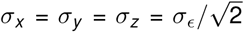 and 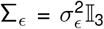. This mathematical model now describes the true looping process over time as well as the resulting noisy measurements.

### Theoretical limits for inferring looping events using simple thresholding

To detect proximity from noisy measurements, we define a detection function *Û* = *ϕ*(*W* ) that maps a measured distance trajectory to an estimated proximity trajectory *Û* = (*û*_1_, …, *û*_*N*_ ). We first focus on a simple threshold-based detection function where *û*_*k*_ = 1 if *d* (*W*_*k*_ ) ≤*α* and *û*_*k*_ = 0 otherwise. In this case, the noisy measurements are used directly as a proxy for the true distance, such that the proximity detection outcome at a time point *k* depends solely on the measurement at that same time point *W*_*k*_ . A potential disadvantage of this approach is that it is agnostic to localization error, which restricts its use to situations where the proximity threshold is large in comparison to the localization error as discussed above. To illustrate this point more quantitatively, we consider an ideal contact where *d* (*V*_*k*_ ) = 0 for all *k*, i.e., the true distance is strictly zero at all times (Fig. **4A**). This represents the bestcase scenario for the purpose of proximity detection, which is useful to discuss fundamental performance limits, without making assumptions about the underlying process *V* . In case of an ideal contact, the measured trajectory *W* is governed entirely by localization error, which is independent across spatial dimensions and time points. To quantify the performance of threshold-based proximity detection, we introduce two measures. The first one is the probability of successful detection *p*_1_ := *P*[*d* (*W*_*k*_ ) ≤ *α* |*d* (*V*_*k*_ ) = 0]. It quantifies for a single time point, how likely the measurement will be within the proximity threshold if the true distance is zero. Since *W*_*k*_ = *ϵ*_*k*_, this probability is given by

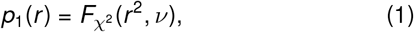

**Figure 4:**
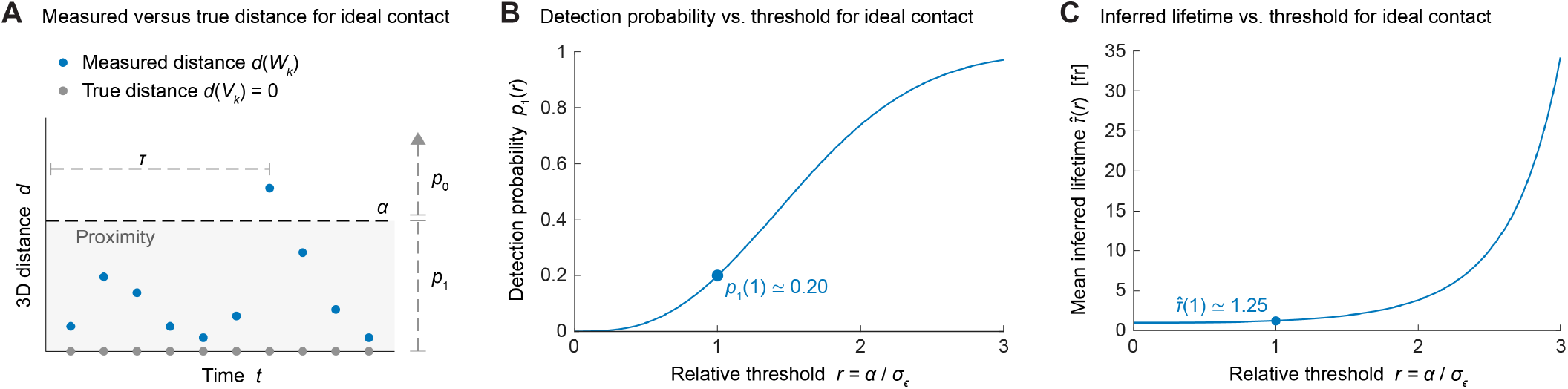
Statistical limits of distance thresholding in the presence of localization error. (A) Illustration of thresholding in the presence of an ideal contact with *d* (*V*_*k*_ ) = 0 for all *k* . Localization error causes the measured distance *d* (*W*_*k*_ ) to be larger than the true distance *d* (*V*_*k*_ ) = 0. The probability of successfully detecting proximity at threshold *α* is denoted by *p*_1_. The inferred interval time *τ* is defined as the first time-point at which the measured distance is larger than the threshold *α*. Its average 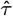 gives an estimate of the typical interval duration that can be resolved before intervals are broken up into shorter pieces. (B) Detection probability as a function of the relative threshold *r* = *α/σ*_*ϵ*_. When the threshold is similar to the distance error *σ*_*ϵ*_ (*r* ≃ 1), the detection probability is *p*_1_ ≃ 0.2. (C) Mean inferred lifetime as a function of the relative threshold *r* = *α/σ*_*ϵ*_. When *r* = *α/σ*_*ϵ*_ = 1, the mean inferred lifetime is 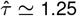 frames.

where 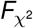 is the cumulative distribution function of a *χ*^2^ random variable with *ν* = 3 degrees of freedom evaluated at *r* ^2^ with *r* = *α/σ*_*ϵ*_ as the normalized proximity threshold. When the threshold *α* is large in comparison to the localization error *σ*_*ϵ*_ (*r* → ∞), the probability *p*_1_ approaches 1. By contrast, if *α* is much smaller than *σ*_*ϵ*_ (*r*→ 0), the probability *p*_1_ vanishes. In the special case where *α* and *σ*_*ϵ*_ are similar (*r*≃ 1), we obtain *p*_1_≃ 0.2 (Fig. **4B**). This means that around 4 out of 5 true proximity events are not detected in this situation.

While the probability of successful detection *p*_1_ captures the overall true positive rate of threshold-based proximity detection, it does not contain information about inferred lifetimes. Because detection functions acting on noisy measurements will eventually break up true proximity intervals into shorter intervals, there exists a maximum recoverable timescale associated to them. To characterize this timescale, we use the mean inferred lifetime 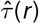 in the case of an ideal, infinitely long contact. To determine this quantity, we calculate the survival function *S*[*k* ] := *P*[*τ > k* |*û*_1_ = 1], which quantifies the probability that greater than *k* time points fall within proximity given that the first time point fell into proximity. Since all measurements are independent across time points, it follows that 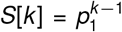 for *k* = 1, …, *N*. Correspondingly, the probability that an inferred proximity event lasts exactly *k* time points can be expressed as 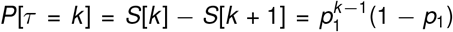. The mean of this distribution is given by

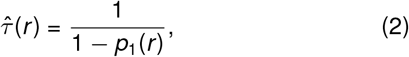

with 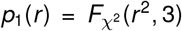. The maximum recoverable timescale starts at 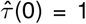 frame and then sharply increases for large *r* (Fig. **4C**). At *r* = 1 we obtain 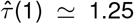 frames, which means that when the localization error *σ*_*ϵ*_ and threshold *α* are comparable, the inferred contact lifetimes will necessarily be around 1 frame, regardless of the true lifetime.

To further illustrate the implications of 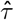, let us consider a sampling rate of *f*_*s*_ = 1 s^*−*1^. To resolve contact lifetimes of up to 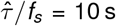 (corresponding to 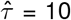 frames) using a proximity threshold *α* = 50 nm, we would require a localization error of approximately *σ*_*ϵ*_ ≃20 nm. If instead the localization error is *σ*_*ϵ*_ = 50 nm, then the proximity threshold cannot be smaller than *α* = 125 nm to resolve proximity intervals of up to 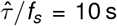. This illustrates a tradeoff between the spatial- and temporal resolution of proximity detection. Another important observation is that the maximum recoverable number of frames of simple thresholding is entirely independent of the sampling rate *f*_*s*_. Therefore, increasing the sampling rate does not allow one to resolve proximity more finely, but instead shifts the inferred lifetimes to shorter values.

### Bayesian thresholding can reduce impact of localization error

We now discuss a potential strategy to overcome the limitations associated with simple thresholding. Specifically, we consider a simple statistical approach, which first deconvolves technical noise from the measured distances, and subsequently performs thresholding on the reconstructed signal. To distinguish signal from noise robustly, we require prior knowledge of the statistics of the measurement noise and the biological signal of interest, which is captured by the covariance matrix of the time series of 3D distance measurements.

We now consider a detection function where *û*_*k*_ = 1 if 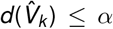 and *û*_*k*_ = 0 otherwise, with 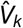 as a statistical reconstruction of the distance *V*_*k*_ from *W*_*k*_ . Here we follow a Bayesian approach and define

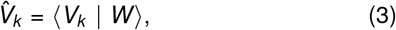

which is the minimum mean-squared error (MMSE) reconstruction of *V*_*k*_ (*84, 85*). In other words, it is the reconstruction that minimizes the expected squared difference to the true distance *V*_*k*_ . A key difference to simple thresholding is that proximity detection is no longer local in time, but instead takes into account information from all time points simultaneously. Correspondingly, the signal reconstruction at time point 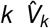 is a function of the entire trajectory *W* . The expectation in (3) is taken with respect to the posterior signal density *p*(*V* |*W* ) ∝ *p*(*V, W* ). If we consider the localization error and the signal to be Gaussian, the posterior density can be obtained in closed form (*84*). For each dimension *i* ∈ {1, 2, 3}, the posterior distribution can be written as 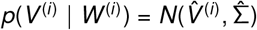, with

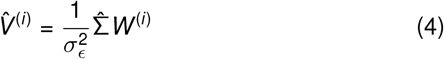

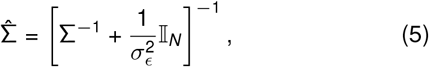

Σ as the *prior* covariance matrix of the signal and I_*N*_ as the identity matrix of size *N* × *N*. Note that since the posterior covariance 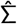 is independent of the measurement *W* ^(*i*)^, and because dimensions are considered isotropic, 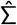 is identical for all dimensions *i* = 1, 2, 3. Once the prior covariance matrix Σ and localization error *σ*_*ϵ*_ have been determined, eqs. (4) and (5) can be computed to obtain the reconstructed distances 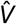, which can then be thresholded to detect proximity intervals.

To illustrate this approach, we performed distance simulations with a prescribed MSD and localization error and compared simple with Bayesian thresholding. We considered sampling intervals of Δ*t* = 12 s and a distance error of 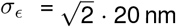. To achieve controlled conditions, the simulations were performed such that the true distance *d* (*V*_*k*_ ) was always larger than a threshold *α* = 50 nm, except for a defined proximity interval of variable length, where *d* (*V*_*k*_ ) was consistently below *α* = 50 nm (Fig. **5A**, top panel). Inferred intervals obtained from simple thresholding are sensitive to distance error, breaking up a single proximity interval into many short ones (Fig. **5A**, middle panel). Deconvolving distance error from measured trajectories using Bayesian methods before thresholding can improve proximity detection substantially (Fig. **5A**, bottom panel).

**Figure 5:**
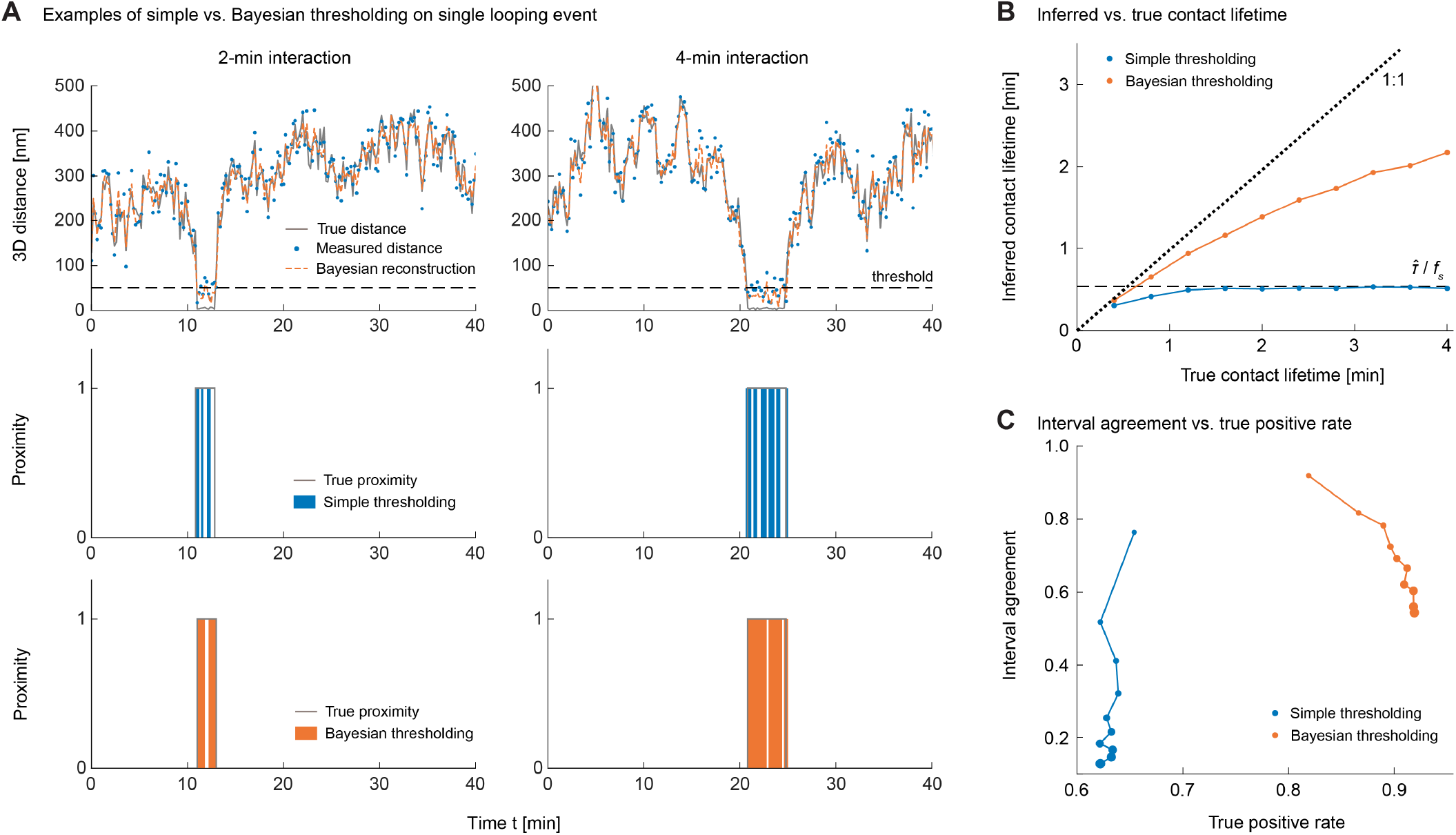
Accounting for localization error in proximity detection using Bayesian thresholding. (A) Totest proximity detection using simple and Bayesian thresholding, we simulated Gaussian distance trajectories with Rouse-like MSD and distance error 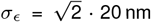. To obtain controlled conditions, the trajectories were simulated such that the true distance was below a threshold of *α* = 50 nm for a defined duration in the middle of the track and above that threshold everywhere else (top panel, black dashed line). Due to distance error, the measured distances can be above or below the threshold at any time point (top panel, blue dots). Bayesian posterior reconstructions of the distance trajectory are shown as dashed orange lines. For inference, the distance covariance matrix Σ was assumed to be known and estimated from the simulated noise-free distance trajectories. Inferred intervals obtained from simple thresholding are shown for two example trajectories in the middle panel (blue rectangles). Results obtained from Bayesian thresholding for the same trajectories are shown in the bottom panel (orange rectangles). (B) Inferred interval length as a function of true interval length. Inferred interval length was calculated as the average inferred interval lifetime obtained via simple thresholding (blue dots) and Bayesian thresholding (orange dots) using *M* = 10000 trajectories each. Averages were computed over all detected intervals across all trajectories for a given condition. Inferred interval lengths from simple thresholding are bounded by the theoretical limit of 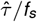 (black dashed line), whereas Bayesian thresholding can resolve timescales beyond that limit. (C) Interval agreement as a function of true positive rate. Interval agreement is defined as *ρ* = 1− |*τ*_inferred_− *τ*_true_ |*/τ*_true_, where *τ*_inferred_ and *τ*_true_ are the average inferred and true interval lengths, respectively. The true positive rate is defined as the probability of correctly detecting proximity, calculated over all time points and trajectories. Dot sizes represent different true interval lengths.

To compare the two approaches more systematically, we determined how the inferred average interval length depends on the true interval length. In line with our earlier considerations, simple thresholding is able to resolve intervals only below the theoretical limit 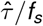, which in this particular case amounts to∼ 0.54 min or equivalently ∼2.7 frames (Fig. **5B**, dashed line). The interval agreement, which we define as *ρ* = 1 −|*τ*_inferred_ −*τ*_true_| */τ*_true_ drops sharply with increasing contact lifetimes, while the true positive rate (i.e., the probability of correctly detecting proximity) remains at approximately 0.6 (Fig. **5C**, blue dots). In the case of Bayesian thresholding, by contrast, the lifetimes can be inferred to a reasonable extent for a larger range of true intervals (Fig. **5B**, orange dots) and the interval agreement *ρ* is substantially less sensitive to the true interval length (Fig. **5C**, orange dots). Likewise, the true positive rate is notably larger across conditions. In summary, these simulations demonstrate that faithful proximity detection requires means to account for and deconvolve localization error from experimental distance measurements.

The simple approach outlined here and the accompanying simulations serve as a proof of principle that the statistical challenges associated with measurement error can be addressed using suitable methods. A full analysis pipeline for real experimental data likely requires coping with additional challenges such as the presence of more stable, longer-lasting proximity events. For example, the Bayesian Inference of Looping Dynamics (BILD) method for detecting CTCF/cohesin-mediated looping events uses a two-state model, in which the looped and unlooped states are calibrated by fitting the MSDs of experimental control conditions (*49, 51*). While BILD can robustly detect loops, it is computationally intensive and requires extensive validation against simulations for hyperparameter selection (*49*). Furthermore, BILD is currently limited to Rouse-like polymer dynamics and needs to be extended to account for MSD exponents different from 0.5 as observed in recent chromatin experiments (*56, 86*).

## Discussion

Live-cell microscopy is ideally suited to study the behavior of chromatin loops over time, and analyses of live-imaging data often seek to estimate the probability, lifetime, and search time of the imaged loops. While 3D distance is the standard criterion by which looping events are identified, great care must be taken to perform the analysis in a rigorous manner and avoid common pitfalls. Most importantly, simple thresholding fails to accurately recover true looping parameters in the presence of even modest localization error. In our analysis of simulated data, with *σ*_*x*_ = 20 nm (which corresponds to a median error on the 3D distance of 26 nm; better than state-of-the-art live-imaging systems (*51, 65, 87, 88*)), thresholding underestimated looping probabilities approximately twofold while misestimating lifetimes and search times in a frame rate-dependent manner. In fact, loop lifetime and search time estimates from thresholding are largely meaningless: these estimates become almost purely a function of the sampling rate in the presence of localization noise. Interestingly, sampling more frequently in time can even worsen lifetime and search time estimates when thresholding is used.

The discrepancy between the true looping parameters and those inferred from thresholding arises entirely from statistical effects. By mathematically formalizing the inference problem, we derived theoretical statistical limits for detecting looping events and inferring their lifetime via simple thresholding. As the uncertainty in 3D distance increases beyond the size of the interaction to be detected, the apparent looping probability approaches zero, and the inferred lifetime approaches a single frame. Recent evidence suggests that functional enhancer-promoter looping interactions occur on scales as small as 25–42 nm (*51*), and live-imaging methods typically have localization errors of at least a few tens of nanometers, so inference is severely affected by thresholding artifacts in this regime.

In the future, live-imaging studies of chromatin looping will greatly benefit from new experimental and computational innovations. While improving spatial and temporal resolution experimentally continues to be an important goal, nonthresholding-based methods must be used to reap its benefits. Bayesian approaches are promising because they can incorporate prior knowledge about the measurement noise and the looping events to be inferred, and information can be integrated across entire trajectories rather than on a pertimepoint basis. Here, we have demonstrated that a simple Bayesian model outperforms thresholding in accurately detecting looping events. The application of Bayesian methods to real data will likely require significant fine-tuning depending on the method of image acquisition and the nature of the looping events to be investigated, and any new inference method must be validated on simulated data with known ground-truth. Equipped with robust statistically validated analysis methods, we expect future studies in this area to contribute towards building an increasingly accurate and precise quantitative bio-physical model of chromatin looping.

## Acknowledgements

The authors thank L. Giorgetti, S. Grosse-Holz, M. K. Huseyin, H. D. Pinholt, and V. Ramanathan for their critical feedback on the manuscript.

## Funding

A.S.H acknowledges funding support from the NIH (DP2GM140938, R33CA257878, R01EB035127, UM1HG011536, R01CA300848, R03OD038390), an NSF CAREER award (2337728), the Gene Regulation Observatory of the Broad Institute of MIT and Harvard, the Novo Nordisk Foundation Center for Genomic Mechanisms of Disease (NNF21SA0072102), a Pew-Stewart Scholar for Cancer Research award, the Mathers Foundation, and an RSC award from the MIT Westaway Fund. This work was supported by the Bridge Project, a partnership between the Koch Institute for Integrative Cancer Research at MIT and the Dana-Farber/Harvard Cancer Center.

## Author contributions

A.S.H. and C.Z. conceived of and designed the project. J.M.J. performed polymer simulation analyses with guidance from A.S.H. C.Z. formulated the statistical treatment and Bayesian thresholding approach. All authors wrote and edited the paper.

## Competing interests

The authors declare no competing interests.

## Data and code availability

The code used to perform the analyses and produce the figures in this work is available at https://github.com/ahansenlab/looping_thresholding_analysis_code.

Large data files are available on Zenodo at https://doi.org/10.5281/zenodo.24096051.

---

**Figure S1:**
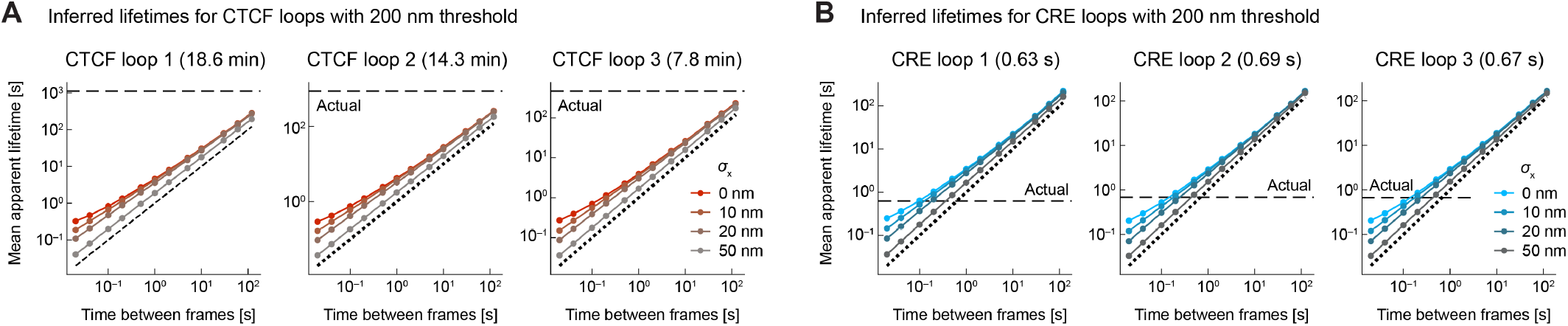
Inference of loop lifetimes with a simple threshold of 200 nm. (A) Same as Fig. **2C** but using a 200-nm threshold: for all three simulated CTCF–CTCF loops, mean apparent lifetimes are shown as a function of the time between frames Δ*t*, for four different noise levels. Dashed line indicate true loop lifetimes and dotted lines indicate mean apparent lifetime = time between frames. (B) Same as (A) but for all three simulated CRE–CRE loops.

**Figure S2:**
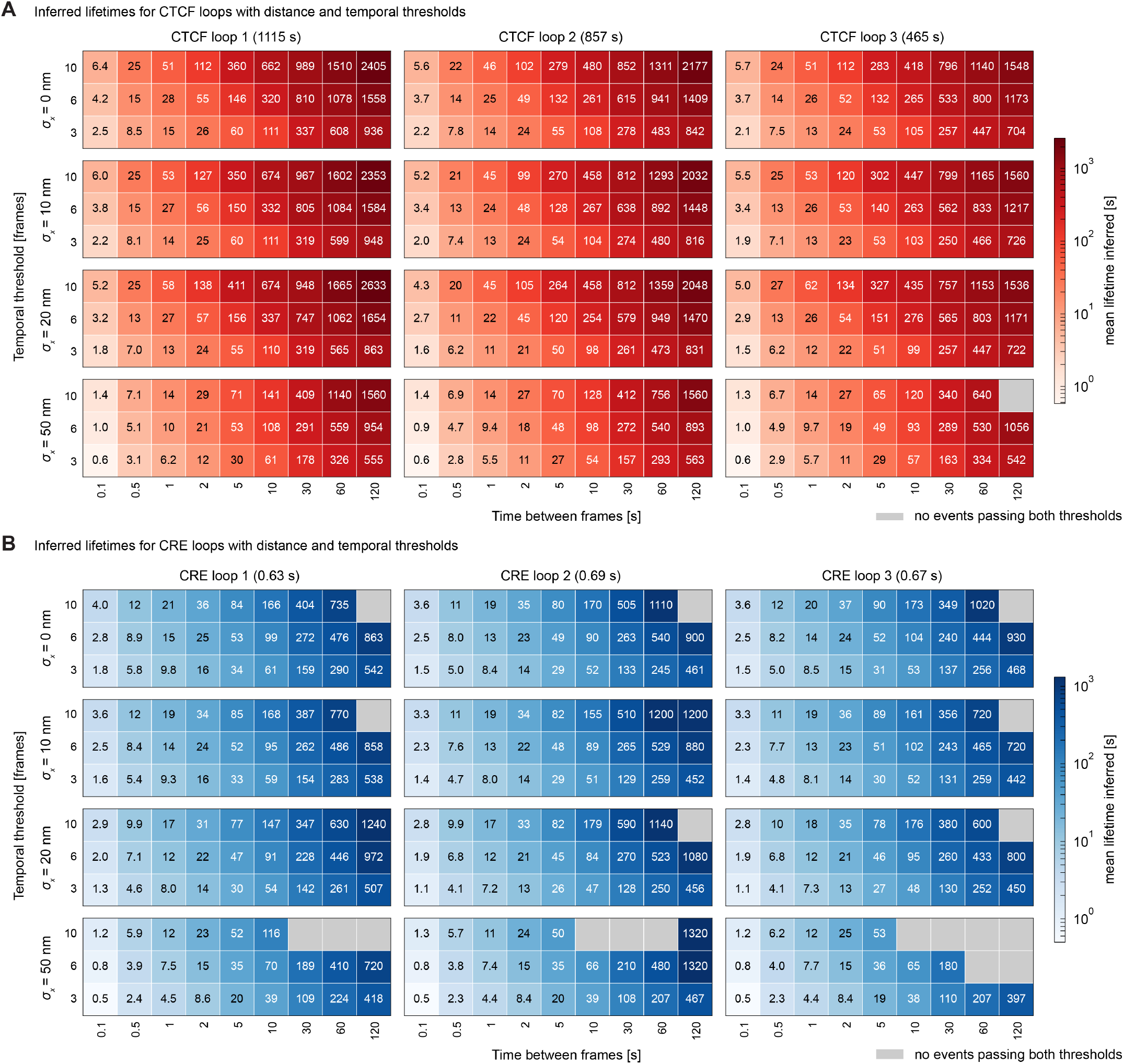
Inference of loop lifetimes with both a distance threshold and temporal threshold. (A) For all three simulated CTCF–CTCF loops and four noise levels, heatmaps of mean inferred lifetime as a function of temporal threshold and time between frames Δ*t* are shown. Individual cells are labeled with the mean inferred lifetime in seconds. Actual lifetimes are indicated in parentheses in plot titles. (B) Same as (A) but for all three simulated CRE–CRE loops.

## Notes

https://github.com/ahansenlab/looping_thresholding_analysis_code

https://doi.org/10.5281/zenodo.24096051

### Competing Interest Statement

The authors have declared no competing interest.

## References

1. J. H. Yang, A. S. Hansen, Enhancer selectivity in space and time: from enhancer–promoter interactions to promoter activation. Nat. Rev. Mol. Cell Biol. 25, 574–591 (2024).

2. C. H. Bohrer, N. A. Fursova, D. R. Larson, Enhancers: A focus on synthetic biology and correlated gene expression. ACS Synth. Biol. 13 (10), 3093–3108 (2024).

3. M. Nollmann, I. Bennabi, M. Götz, T. Gregor, The impact of space and time on the functional output of the genome. Cold Spring Harb. Perspect. Biol. 14 (4), a040378 (2022).

4. J. Dekker, L. A. Mirny, The chromosome folding problem and how cells solve it. Cell 187 (23), 6424–6450 (2024).

5. A. Marin-Gonzalez, T. Ha, Folding a broken genome: the versatile roles of cohesin in genome maintenance. Nat. Rev. Genet. 27, 762–778 (2026).

6. J.-M. Peters, How DNA loop extrusion mediated by cohesin enables V(D)J recombination. Curr. Opin. Cell Biol. 70, 75–83 (2021).

7. N. Chen, S. C. B. Buonomo, Three-dimensional nuclear organisation and the DNA replication timing program. Curr. Opin. Struct. Biol. 83, 102704 (2023).

8. E. Lieberman-Aiden, N. L. van Berkum, L. Williams, M. Imakaev, T. Ragoczy, A. Telling, I. Amit, B. R. Lajoie, P. J. Sabo, M. O. Dorschner, R. Sandstrom, B. Bernstein, M. A. Bender, M. Groudine, A. Gnirke, J. Stamatoyannopoulos, L. A. Mirny, E. S. Lander, J. Dekker, Comprehensive mapping of long-range interactions reveals folding principles of the human genome. Science 326 (5950), 289–293 (2009).

9. J. M. Jusuf, J. H. Yang, J. Toppen, S. Grosse-Holz, M. Gabriele, P. Mach, I. M. Flyamer, C. Zechner, L. Giorgetti, L. A. Mirny, A. S. Hansen, Genome-wide absolute quantification of chromatin looping. Nat. Struct. Mol. Biol. 33 (7), 1105–1114 (2026).

10. S. S. Rao, M. H. Huntley, N. C. Durand, E. K. Stamenova, I. D. Bochkov, J. T. Robinson, A. L. Sanborn, I. Machol, A. D. Omer, E. S. Lander, E. L. Aiden, A 3D map of the human genome at kilobase resolution reveals principles of chromatin looping. Cell 159 (7), 1665–1680 (2014).

11. A. S. Hansen, CTCF as a boundary factor for cohesinmediated loop extrusion: evidence for a multi-step mechanism. Nucleus 11 (1), 132–148 (2020).

12. Y. Kim, Z. Shi, H. Zhang, I. J. Finkelstein, H. Yu, Human cohesin compacts DNA by loop extrusion. Science 366 (6471), 1345–1349 (2019).

13. I. F. Davidson, B. Bauer, D. Goetz, W. Tang, G. Wutz, J.-M. Peters, DNA loop extrusion by human cohesin. Science 366 (6471), 1338–1345 (2019).

14. J. R. Dixon, S. Selvaraj, F. Yue, A. Kim, Y. Li, Y. Shen, M. Hu, J. S. Liu, B. Ren, Topological domains in mammalian genomes identified by analysis of chromatin interactions. Nature 485 (7398), 376–380 (2012).

15. E. P. Nora, B. R. Lajoie, E. G. Schulz, L. Giorgetti, I. Okamoto, N. Servant, T. Piolot, N. L. van Berkum, J. Meisig, J. Sedat, J. Gribnau, E. Barillot, N. Blüthgen, J. Dekker, E. Heard, Spatial partitioning of the regulatory landscape of the X-inactivation centre. Nature 485 (7398), 381–385 (2012).

16. J. A. Beagan, J. E. Phillips-Cremins, On the existence and functionality of topologically associating domains. Nat. Genet. 52 (1), 8–16 (2020).

17. K. L. Kim, G. J. Rahme, V. Y. Goel, C. A. E. Farran, A. S. Hansen, B. E. Bernstein, Dissection of a CTCF topological boundary uncovers principles of enhancer-oncogene regulation. Mol. Cell 84 (7), 1365–1376.e7 (2024).

18. S. Chakraborty, N. Wenzlitschke, M. J. Anderson, A. Eraso, M. Baudic, J. J. Thompson, A. A. Evans, L. M. Shatford-Adams, R. Chari, P. Awasthi, R. K. Dale, M. Lewandoski, T. J. Petros, P. P. Rocha, Deletion of a single CTCF motif at the boundary of a chromatin domain with three FGF genes disrupts gene expression and embryonic development. Dev. Cell 60, 1838–1853.e9 (2025).

19. D.G. Lupiáñez, K. Kraft, V. Heinrich, P. Krawitz, F. Brancati, E. Klopocki, D. Horn, H. Kayserili, J. M. Opitz, R. Laxova, F. Santos-Simarro, B. Gilbert-Dussardier, L. Wittler, M. Borschiwer, S. A. Haas, M. Osterwalder, M. Franke, B. Timmermann, J. Hecht, M. Spielmann, A. Visel, S. Mundlos, Disruptions of topological chromatin domains cause pathogenic rewiring of gene-enhancer interactions. Cell 161 (5), 1012–1025 (2015).

20. O. Symmons, V. V. Uslu, T. Tsujimura, S. Ruf, S. Nassari, W. Schwarzer, L. Ettwiller, F. Spitz, Functional and topological characteristics of mammalian regulatory domains. Genome Res. 24 (3), 390–400 (2014).

21. J. H. Yang, H. B. Brandão, A. S. Hansen, DNA double-strand break end synapsis by DNA loop extrusion. Nat. Commun. 14 (1), 1913 (2023).

22. C. Arnould, V. Rocher, A.-L. Finoux, T. Clouaire, K. Li, F. Zhou, P. Caron, P. E. Mangeot, E. P. Ricci, R. Mourad, J. E. Haber, D. Noordermeer, G. Legube, Loop extrusion as a mechanism for formation of DNA damage repair foci. Nature 590 (7847), 660–665 (2021).

23. Y. Liu, R. S. Zou, S. He, Y. Nihongaki, X. Li, S. Razavi, B. Wu, T. Ha, Very fast CRISPR on demand. Science 368 (6496), 1265–1269 (2020).

24. Z. Ba, J. Lou, A. Y. Ye, H.-Q. Dai, E. W. Dring, S. G. Lin, S. Jain, N. Kyritsis, K.-R. Kieffer-Kwon, R. Casellas, F. W. Alt, CTCF orchestrates long-range cohesin-driven V(D)J recombinational scanning. Nature 586 (7828), 305–310 (2020).

25. D. Bsteh, H. F. Moussa, G. Michlits, R. Yelagandula, J. Wang, U. Elling, O. Bell, Loss of cohesin regulator PDS5A reveals repressive role of polycomb loops. Nat. Commun. 14 (1), 8160 (2023).

26. Y. Ogiyama, B. Schuettengruber, G. L. Papadopoulos, J.-M. Chang, G. Cavalli, Polycomb-dependent chromatin looping contributes to gene silencing during Drosophila development. Mol. Cell 71 (1), 73–88.e5 (2018).

27. K. Kraft, K. E. Yost, S. E. Murphy, A. Magg, Y. Long, M. R. Corces, J. M. Granja, L. Wittler, S. Mundlos, T. R. Cech, A. N. Boettiger, H. Y. Chang, Polycomb-mediated genome architecture enables long-range spreading of H3K27 methylation. Proc. Natl. Acad. Sci. U. S. A. 119 (22), e2201883119 (2022).

28. S. Kim, M. K. Huseyin, A. S. Hansen, Polycomb-mediated 3D chromatin interactions in gene regulation. Curr. Opin. Genet. Dev. 97, 102448 (2026).

29. B. Pang, M. P. Snyder, Systematic identification of silencers in human cells. Nat. Genet. 52 (3), 254–263 (2020).

30. J. A. Segert, S. S. Gisselbrecht, M. L. Bulyk, Transcriptional silencers: Driving gene expression with the brakes on. Trends Genet. 37 (6), 514–527 (2021).

31. X. Wang, F. Yue, Hijacked enhancer-promoter and silencer-promoter loops in cancer. Curr. Opin. Genet. Dev. 86, 102199 (2024).

32. X. Li, M. Levine, What are tethering elements? Curr. Opin. Genet. Dev. 84, 102151 (2024).

33. G. Mohana, J. Dorier, X. Li, M. Mouginot, R. C. Smith, H. Malek, M. Leleu, D. Rodriguez, J. Khadka, P. Rosa, P. Cousin, C. Iseli, S. Restrepo, N. Guex, B. D. McCabe, A. Jankowski, M. S. Levine, M. C. Gambetta, Chromosome-level organization of the regulatory genome in the Drosophila nervous system. Cell 186 (18), 3826– 3844.e26 (2023).

34. G. Bower, E. W. Hollingsworth, S. H. Jacinto, J. A. Alcantara, B. Clock, K. Cao, M. Liu, A. Dziulko, A. Alcaina-Caro, Q. Xu, D. Skowronska-Krawczyk, J. Lopez-Rios, D. E. Dickel, A. F. Bardet, L. A. Pennacchio, A. Visel, E. Z. Kvon, Range extender mediates long-distance enhancer activity. Nature 643, 830–838 (2025).

35. J. W. Blayney, H. Francis, A. Rampasekova, B. Camellato, L. Mitchell, R. Stolper, L. Cornell, C. Babbs, J. D. Boeke, D. R. Higgs, M. Kassouf, Super-enhancers include classical enhancers and facilitators to fully activate gene expression. Cell 186 (26), 5826–5839.e18 (2023).

36. V. Ramanathan, C.-J. Guo, M. Nagano, V. Y. Goel, C. K. Y. Hong, A. D. Swett, A. Caulier, E. King, Z. Tothova, V. G. Sankaran, A. S. Hansen, Cohesin loading at regulatory elements shapes 3D genome folding during erythropoiesis. bioRxiv (2026). doi: 10.64898/2026.08.02.741886.

37. T.-H. S. Hsieh, C. Cattoglio, E. Slobodyanyuk, A. S. Hansen, O. J. Rando, R. Tjian, X. Darzacq, Resolving the 3D landscape of transcription-linked mammalian chromatin folding. Mol. Cell 78 (3), 539–553.e8 (2020).

38. N. Krietenstein, S. Abraham, S. V. Venev, N. Abdennur, J. Gibcus, T.-H. S. Hsieh, K. M. Parsi, L. Yang, R. Maehr, L. A. Mirny, J. Dekker, O. J. Rando, Ultrastructural details of mammalian chromosome architecture. Mol. Cell 78 (3), 554–565.e7 (2020).

39. C. K. Hong, F. Feng, V. Ramanathan, J. Liu, A. S. Hansen, Genome structure mapping with high-resolution 3D genomics and deep learning. bioRxiv (2025). doi: 10.1101/2025.05.06.650874.

40. J. Dekker, F. Alber, S. Aufmkolk, B. J. Beliveau, B. G. Bruneau, A. S. Belmont, L. Bintu, A. Boettiger, R. Calandrelli, C. M. Disteche, et al., Spatial and temporal organization of the genome: current state and future aims of the 4D nucleome project. Mol. Cell 83 (15), 2624–2640 (2023).

41. B. Akgol Oksuz, L. Yang, S. Abraham, S. V. Venev, N. Krietenstein, K. M. Parsi, H. Ozadam, M. E. Oomen, A. Nand, H. Mao, R. M. J. Genga, R. Maehr, O. J. Rando, L. A. Mirny, J. H. Gibcus, J. Dekker, Systematic evaluation of chromosome conformation capture assays. Nat. Methods 18 (9), 1046–1055 (2021).

42. T.-H. S. Hsieh, A. Weiner, B. Lajoie, J. Dekker, N. Friedman, O. J. Rando, Mapping nucleosome resolution chromosome folding in yeast by Micro-C. Cell 162 (1), 108– 119 (2015).

43. A. S. Hansen, T.-H. S. Hsieh, C. Cattoglio, I. Pustova, R. Saldaña-Meyer, D. Reinberg, X. Darzacq, R. Tjian, Distinct classes of chromatin loops revealed by deletion of an RNA-binding region in CTCF. Mol. Cell 76 (3), 395– 411.e13 (2019).

44. T. Yang, S. Wang, Image-based 3D genomics through chromatin tracing. Nat. Rev. Methods Primers 4 (1), 1–21 (2024).

45. Y. Takei, Y. Yang, J. White, I. N. Goronzy, J. Yun, M. Prasad, L. J. Ombelets, S. Schindler, P. Bhat, M. Guttman, L. Cai, Spatial multi-omics reveals cell-typespecific nuclear compartments. Nature 641 (8064), 1037– 1047 (2025).

46. J.-H. Su, P. Zheng, S. S. Kinrot, B. Bintu, X. Zhuang, Genome-scale imaging of the 3D organization and transcriptional activity of chromatin. Cell 182 (6), 1641– 1659.e26 (2020).

47. A. S. Labade, Z. D. Chiang, C. Comenho, P. L. Reginato, A. C. Payne, A. S. Earl, R. Shrestha, F. M. Duarte, E. Habibi, R. Zhang, G. M. Church, E. S. Boyden, F. Chen, J. D. Buenrostro, Expansion in situ genome sequencing links nuclear abnormalities to aberrant chromatin regulation. Science 389, eadt2781 (2025).

48. H. B. Brandão, M. Gabriele, A. S. Hansen, Tracking and interpreting long-range chromatin interactions with superresolution live-cell imaging. Curr. Opin. Cell Biol. 70, 18– 26 (2021).

49. M. Gabriele, H. B. Brandão, S. Grosse-Holz, A. Jha, G. Dailey, C. Cattoglio, T.-H. S. Hsieh, L. A. Mirny, C. Zechner, A. S. Hansen, Dynamics of CTCF-and cohesin-mediated chromatin looping revealed by live-cell imaging. Science 376 (6592), 496–501 (2022).

50. P. Mach, P. I. Kos, Y. Zhan, J. Cramard, S. Gaudin, J. Tünnermann, E. Marchi, J. Eglinger, J. Zuin, M. Kryzhanovska, S. Smallwood, L. Gelman, G. Roth, E. P. Nora, G. Tiana, L. Giorgetti, Cohesin and CTCF control the dynamics of chromosome folding. Nat. Genet. 54 (12), 1907–1918 (2022).

51. J. H. Yang, H. D. Pinholt, J. Toppen, M. K. Huseyin, J. M. Jusuf, C. C. Katsifis, J. Kæstel-Hansen, L. A. Mirny, C. Zechner, A. S. Hansen, Live-cell imaging of enhancer-promoter dynamics reveals transient contact-driven gene activation. bioRxiv (2026). doi: 10.64898/2026.06.21.733446.

52. T. Sabaté, B. Lelandais, M.-C. Robert, M. Szalay, J.-Y. Tinevez, E. Bertrand, C. Zimmer, Uniform dynamics of cohesin-mediated loop extrusion in living human cells. Nat. Genet. 57 (12), 3152–3164 (2025).

53. H. Chen, M. Levo, L. Barinov, M. Fujioka, J. B. Jaynes, T. Gregor, Dynamic interplay between enhancer–promoter topology and gene activity. Nat. Genet. 50 (9), 1296–1303 (2018).

54. L. Cheng, C. De, J. Li, R. Korff, A. Pertsinidis, Mechanisms of transcription control by distal enhancers from single-gene nanoscopy. Cell pp. S0092–8674(26)01130– X (2026).

55. K. Maeshima, The shifting paradigm of chromatin structure: from the 30-nm chromatin fiber to liquid-like organization. Proc. Jpn. Acad. Ser. B Phys. Biol. Sci. 101 (6), 339–356 (2025).

56. M. Mazzocca, D. N. Narducci, S. Grosse-Holz, J. Matthias, T. S. Karpova, A. S. Hansen, Integrated MINFLUX tracking reveals two distinct chromatin dynamics classes across cell types. Nat. Struct. Mol. Biol. 33 (6), 905–914 (2026).

57. Y. Zhu, W. E. Moerner, L. S. Qi, CRISPR-Cas-based live cell imaging of genome dynamics. Nat. Rev. Genet. 27, 622––641 (2026).

58. J. M. Alexander, J. Guan, B. Li, L. Maliskova, M. Song, Y. Shen, B. Huang, S. Lomvardas, O. D. Weiner, Livecell imaging reveals enhancer-dependent Sox2 transcription in the absence of enhancer proximity. eLife 8, e41769 (2019).

59. J. Rodriguez, G. Ren, C. R. Day, K. Zhao, C. C. Chow, D. R. Larson, Intrinsic dynamics of a human gene reveal the basis of expression heterogeneity. Cell 176 (1-2), 213–226.e18 (2019).

60. Y. Wan, D. G. Anastasakis, J. Rodriguez, M. Palangat, P. Gudla, G. Zaki, M. Tandon, G. Pegoraro, C. C. Chow, M. Hafner, D. R. Larson, Dynamic imaging of nascent RNA reveals general principles of transcription dynamics and stochastic splice site selection. Cell 184 (11), 2878– 2895.e20 (2021).

61. W. J. de Jonge, H. P. Patel, J. V. W. Meeussen, T. L. Lenstra, Following the tracks: How transcription factor binding dynamics control transcription. Biophys. J. 121 (9), 1583–1592 (2022).

62. E. Soutoglou, J. F. Dorn, K. Sengupta, M. Jasin, A. Nussenzweig, T. Ried, G. Danuser, T. Misteli, Positional stability of single double-strand breaks in mammalian cells. Nat. Cell Biol. 9 (6), 675–682 (2007).

63. M. Du, F. Zou, Y. Li, Y. Yan, L. Bai, Chemically induced chromosomal interaction (CICI) method to study chromosome dynamics and its biological roles. Nat. Commun. 13 (1), 757 (2022).

64. F. Zou, Y. Li, T. Földes, H. D. Pinholt, C. Smith, L. Mirny, L. Bai, Condensin accelerates long-range intrachromosomal interactions. Nat. Commun. 17 (2026).

65. M. Ubertini, N. Louafi, K. Landry, P. I. Kos, G. Roth, E. Marchi, J. Cramard, G. Tiana, L. Giorgetti, Loop extrusion creates rare, long-lived encounters underlying enhancer-promoter communication. bioRxiv (2025). doi: 10.1101/2025.09.24.678119.

66. D. Mathog, M. Hochstrasser, Y. Gruenbaum, H. Saumweber, J. Sedat, Characteristic folding pattern of polytene chromosomes in Drosophila salivary gland nuclei. Nature 308 (5958), 414–421 (1984).

67. M. Hochstrasser, D. Mathog, Y. Gruenbaum, H. Saumweber, J. W. Sedat, Spatial organization of chromosomes in the salivary gland nuclei of Drosophila melanogaster. J. Cell Biol. 102 (1), 112–123 (1986).

68. B. Trask, D. Pinkel, G. van den Engh, The proximity of DNA sequences in interphase cell nuclei is correlated to genomic distance and permits ordering of cosmids spanning 250 kilobase pairs. Genomics 5 (4), 710–717 (1989).

69. M. Lakadamyali, M. P. Cosma, Visualizing the genome in high resolution challenges our textbook understanding. Nat. Methods 17 (4), 371–379 (2020).

70. R. E. Thompson, D. R. Larson, W. W. Webb, Precise nanometer localization analysis for individual fluorescent probes. Biophys. J. 82 (5), 2775–2783 (2002).

71. Y. Gao, M. L. Kilfoil, Accurate detection and complete tracking of large populations of features in three dimensions. Opt. Express 17 (6), 4685–4704 (2009).

72. A. Dominguez Mantes, A. Herrera, I. Khven, A. Schlaeppi, E. Kyriacou, G. Tsissios, E. Skoufa, L. Santangeli, E. Buglakova, E. B. Durmus, S. Manley, A. Kreshuk, D. Arendt, C. Aztekin, J. Lingner, G. La Manno, M. Weigert, Spotiflow: accurate and efficient spot detection for fluorescence microscopy with deep stereographic flow regression. Nat. Methods 22 (7), 1495–1504 (2025).

73. S. C. M. Reinhardt, L. A. Masullo, I. Baudrexel, P. R. Steen, R. Kowalewski, A. S. Eklund, S. Strauss, E. M. Unterauer, T. Schlichthaerle, M. T. Strauss, C. Klein, R. Jungmann, Ångström-resolution fluorescence microscopy. Nature 617 (7962), 711–716 (2023).

74. J. Lee, L.-F. Chen, S. Gaudin, K. Gupta, A. Spakowitz, A. N. Boettiger, Kinetic organization of the genome revealed by ultra-resolution, multiscale live imaging. bioRxiv (2025). doi: 10.1101/2025.03.27.645817.

75. A. Hafner, M. Park, S. E. Berger, S. E. Murphy, E. P. Nora, A. N. Boettiger, Loop stacking organizes genome folding from TADs to chromosomes. Mol. Cell 83 (9), 1377– 1392.e6 (2023).

76. T. Sabaté, B. Lelandais, E. Bertrand, C. Zimmer, Polymer simulations guide the detection and quantification of chromatin loop extrusion by imaging. Nucleic Acids Res. 51 (6), 2614–2632 (2023).

77. N. Khanna, Y. Zhang, J. S. Lucas, O. K. Dudko, C. Murre, Chromosome dynamics near the sol-gel phase transition dictate the timing of remote genomic interactions. Nat. Commun. 10 (1), 2771 (2019).

78. J. S. Lucas, Y. Zhang, O. K. Dudko, C. Murre, 3D trajectories adopted by coding and regulatory DNA elements: first-passage times for genomic interactions. Cell 158 (2), 339–352 (2014).

79. A. Amitai, D. Holcman, Analysis of chromatin dynamics and search processes in the nucleus, in Modeling the 3D Conformation of Genomes (CRC Press, Boca Raton, FL), pp. 177–206 (2019).

80. A. Amitai, A. Seeber, S. M. Gasser, D. Holcman, Visualization of chromatin decompaction and break site extrusion as predicted by statistical polymer modeling of singlelocus trajectories. Cell Rep. 18 (5), 1200–1214 (2017).

81. J.-M. Arbona, S. Herbert, E. Fabre, C. Zimmer, Inferring the physical properties of yeast chromatin through Bayesian analysis of whole nucleus simulations. Genome Biol. 18 (1), 81 (2017).

82. V. I. P. Keizer, S. Grosse-Holz, M. Woringer, L. Zambon, K. Aizel, M. Bongaerts, F. Delille, L. Kolar-Znika, V. F. Scolari, S. Hoffmann, E. J. Banigan, L. A. Mirny, M. Dahan, D. Fachinetti, A. Coulon, Live-cell micromanipulation of a genomic locus reveals interphase chromatin mechanics. Science 377 (6605), 489–495 (2022).

83. M. P. H. Stumpf, M. A. Porter, Critical truths about power laws. Science 335 (6069), 665–666 (2012).

84. C. M. Bishop, Pattern recognition and machine learning (Springer, New York, NY) (2006).

85. S. M. Kay, Fundamentals of Statistical Signal Processing, Volume I: Estimation Theory, Prentice-Hall Signal Processing Series (Prentice Hall PTR, Upper Saddle River, NJ) (1993).

86. S. Grosse-Holz, L. A. Mirny, A. Coulon, Scale-free models of chromosome structure, dynamics, and mechanics. PNAS Nexus 4 (12), pgaf337 (2025).

87. Y. Zhu, A. Balaji, M. Han, L. Andronov, A. R. Roy, Z. Wei, C. Chen, L. Miles, S. Cai, Z. Gu, A. Tse, B. C. Yu, T. Uenaka, X. Lin, A. J. Spakowitz, W. E. Moerner, L. S. Qi, High-resolution dynamic imaging of chromatin DNA communication using Oligo-LiveFISH. Cell 188 (12), 3310– 3328.e27 (2025).

88. A. Raterink, R. P. Ghosh, L. Yang, Q. Shi, M. K. Nguyen, I. B. Hilton, J. T. Liphardt, A.-K. Gustavsson, Robust fluorescent labeling and tracking of endogenous non-repetitive genomic loci. bioRxiv (2025). doi: 10.1101/2025.08.22.671818.

